# Semi-automated identification of southern right whales from drone imagery by classical image matching methods

**DOI:** 10.64898/2026.07.19.739396

**Authors:** Ben Fabry, Greta Jacobs, Johannes Bartl, Max Fabry

## Abstract

Individual southern right whales (*Eubalaena australis*) can be identified from the callosity pattern on the head, a stable pattern of roughened keratinized patches. Manual comparison of drone images with large catalogs, however, is time-consuming. For southern right whales, automated photo-identification approaches typically require substantial training data or retraining when new individuals are added. Here we evaluate two classical image-similarity methods for individual identification from standardized dorsal head images: histograms of oriented gradients (HOG), and symmetric log-chamfer distance. We tested 375 query images from 198 known whales with recurrent sightings against 411 reference images, one for each individual whale. HOG ranked the correct whale first in 361 of 375 cases (96.3%), whereas log-chamfer ranked the correct whale first in 355 cases (94.7%). All incorrect rank-1 matches could be identified by a high risk score computed from query-reference distance distributions. A combined rule selecting the first-ranked candidate from the method with the lower risk score increased rank-1 accuracy to 368 cases (98.1%). These results show that classical registered image matching provides a practical tool for southern right whale photo-identification.

## Introduction

Long-term monitoring of southern right whales (*Eubalaena australis*) depends on the ability to recognize individuals across repeated sightings. The main natural mark used for individual recognition is the callosity pattern on the head, including the bonnet, rostral islands, lip patches, coaming, and post blow-hole island. These keratinized patches are colonized by cyamids and appear as pale markings against the dark skin. They produce persistent individual patterns that can be photographed from aircraft, vessels, shore vantage points, and drones (Best 1990, Hiby & Lovell 2001).

Photo-identification from callosity patterns can be applied retrospectively to historical catalogs, and provides the individual encounter histories needed for estimating survival, calving intervals, site fidelity, abundance, movement, habitat use, health, and reproductive history (Bannister 2001, Best 1990, Bogucki *et al*. 2019, Khan *et al*. 2022). Established regional catalogs such as the Australasian Right Whale Photo-Identification Catalogue use these records to support mark-recapture analyses, conservation assessment, and management (Khan et al. 2022). However, manual matching is labor-intensive. A new image may need to be compared against hundreds or thousands of catalog images, and matching performance can be affected by image angle, whale posture, partial surfacing, glare, calf callosity development, annotation quality, and observer experience.

Several approaches have been developed to support and speed up manual matching. One early computer-assisted system converted manually outlined callosity patches into simplified shape- and-position descriptors and compared them across photographs to generate a shortlist of likely matches, which greatly reduced the number of by-eye comparisons (Hiby & Lovell 2001). More recently, deep-learning workflows have been applied to right whale photo-identification, including head detection, image standardization, and individual classification with convolutional neural networks (CNNs). For North Atlantic right whales, a CNN-based classifier achieved approximately 87% rank-1 accuracy when matching images to a catalog of 447 known individuals (Bogucki et al. 2019, Khan et al. 2022). At a broader cetacean scale, the open-source photo-identification platform Flukebook provides infrastructure for whale and dolphin photo-identification by combining image archives, sighting records, and automated matching algorithms. Its database contains more than 2 million photographs and supports feature-based matching, contour-based matching, and learned image representations (Blount *et al*. 2022, Patton *et al*. 2023).

These systems are powerful, but their practical reliability depends on training-set composition, catalog curation, image quality, and how easily new identities can be added. This is particularly important for southern right whales, where catalogs may contain many individuals with only a small number of re-sightings and where image appearance varies with viewpoint, surfacing posture, sea state, and callosity development (Khan et al. 2022).

Closed-set classifier-based systems can identify only individuals represented in the training set, and adding new individuals typically requires retraining. Open-set image-similarity approaches, such as HotSpotter/SIFT-based matching, CurvRank, or deep metric-learning embeddings, reduce this limitation by comparing a new image with catalog images rather than assigning it directly to a fixed identity class. For southern right whales, however, automated matching remains challenging: in a multi-species benchmark, southern right whales were among the least accurately recognized species (Patton et al. 2023). Another study that used pose invariant embeddings to match lateral head photographs reported that the correct whale was ranked first in only 16% of cases (Khan et al. 2022). Some of these approaches, particularly learned contour or image-representation models, still require representative training data, and all require careful human verification (Blount et al. 2022, Khan et al. 2022). This leaves a practical need for methods that are accurate, easy to adapt and implement, and not dependent on repeated training examples for each individual.

Here, we evaluate two classical open-set image-similarity approaches for southern right whale identification from segmented callosity masks: Histogram of Oriented Gradients (HOG) descriptor distance, and symmetric log-chamfer edge distance. These simple image matching methods are model-free, do not require training, and function with only a single catalog image. The purpose of this paper is to describe the workflow and registration procedure, compare the performance of the two methods, and report their usefulness for identifying individual whales in a regional southern right whale catalog.

## Methods

### Field surveys and image acquisition

Images were collected during drone-based surveys of southern right whales along the south-western Australian coast between 2022 and 2024. Animal ethics approval was granted through the Animal Ethics Committee of the University of Western Australia (project 2021/ET000236). Surveys were conducted during the calving season from shore-accessible aggregation areas. Whales were located either by observers scanning from elevated shore positions or during manual drone transects along low-lying beaches. When a whale was detected, a DJI Phantom 4 Pro drone (DJI, Shenzhen, China) equipped with a 1-inch 20 MP CMOS sensor and an 8.8 mm lens (24 mm equivalent focal length; 84° FOV; aperture f/2.8-f/11) was positioned above the animal, typically at approximately 20 m altitude. Images or video frames were collected when the whale surfaced. These images were used for individual identification from the head callosity pattern.

### Reference database and segmentation

A total of 1,660 images were curated into a catalog and database in which each identified whale is associated with one passport photo-like reference image. A reference image was selected from all available images of an identified whale based on the quality with which the callosity pattern can be seen, ideally from a near-vertical perspective. Reference images were manually rotated so that heads point left, shifted so the midline of the head aligns with the horizontal center line, zoomed and cropped so that the bonnet and post blow-hole islands fall at defined image positions (Fig. 1), and resized to standardized 1200 x 1000 pixel images. The callosity pattern was then manually segmented as a binary mask. Matching was restricted to the central head region of interest (ROI) with x = 225 to 975 and y = 350 to 650 (corresponding to a 750 x 300 pixel crop) in the standardized mask (Fig. 1). This region captures the central dorsal callosity pattern while excluding the eye patches and parts of the bonnet that are typically below the water surface.

**Fig. 1.**
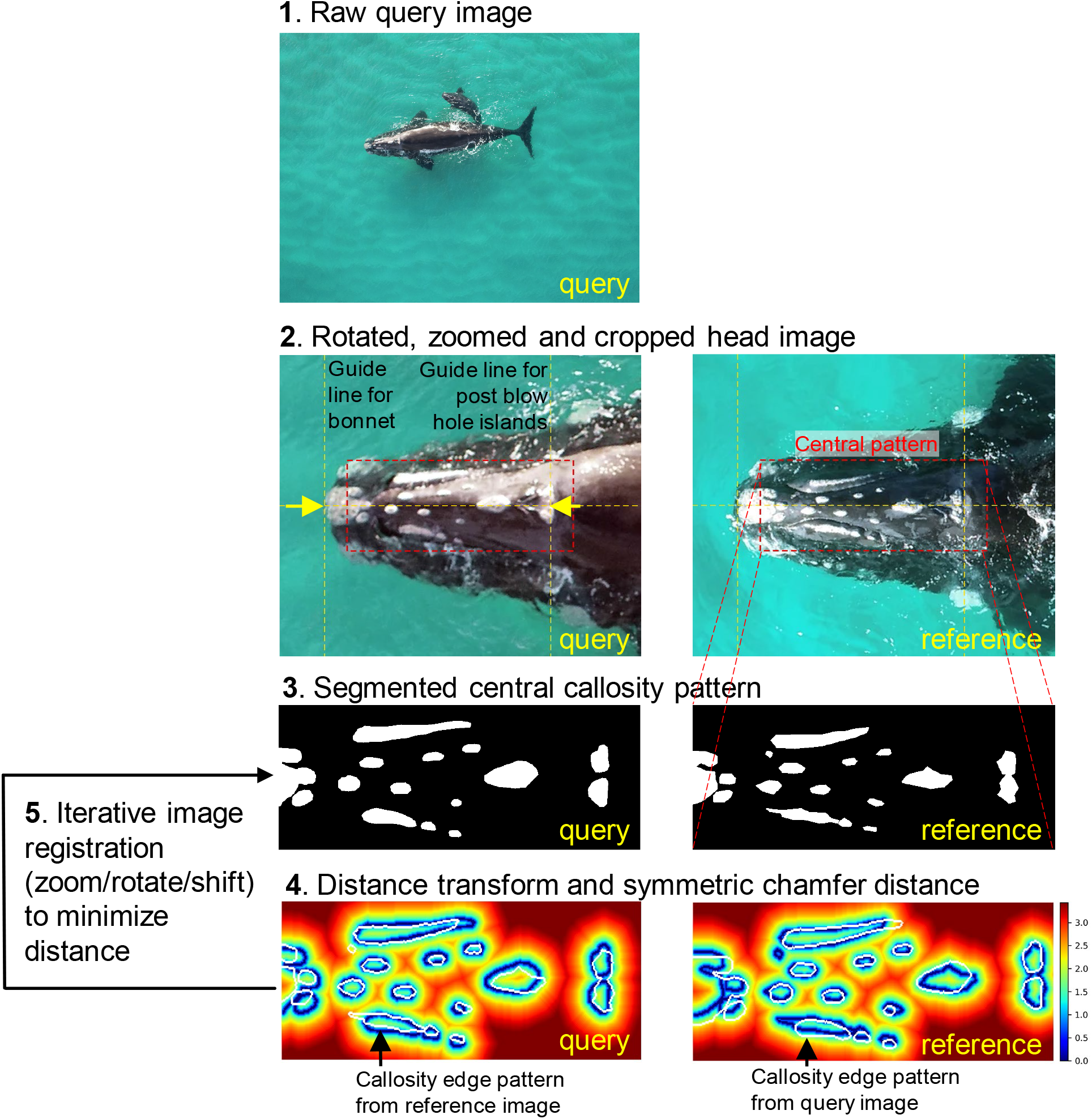
Workflow for callosity-based image matching. A raw drone image was converted into a standardized query gallery crop. Yellow dashed guide lines indicate the fixed alignment lines used during gallery curation, and the red dashed rectangle marks the region of interest (ROI) used for matching. The reference image was processed in the same coordinate system. The query and reference callosity patterns were manually segmented as binary masks. For chamfer matching, callosity edges were extracted from each full mask before ROI cropping and converted into log-distance images, log(pixel distance + 1), after capping distances at 30 pixels. The opposite mask edge was then transformed by the best registration and overlaid on the distance image. The HOG distance method uses the same full-mask-before-crop edge extraction principle to avoid ROI-boundary artefacts.

**Fig. 2.**
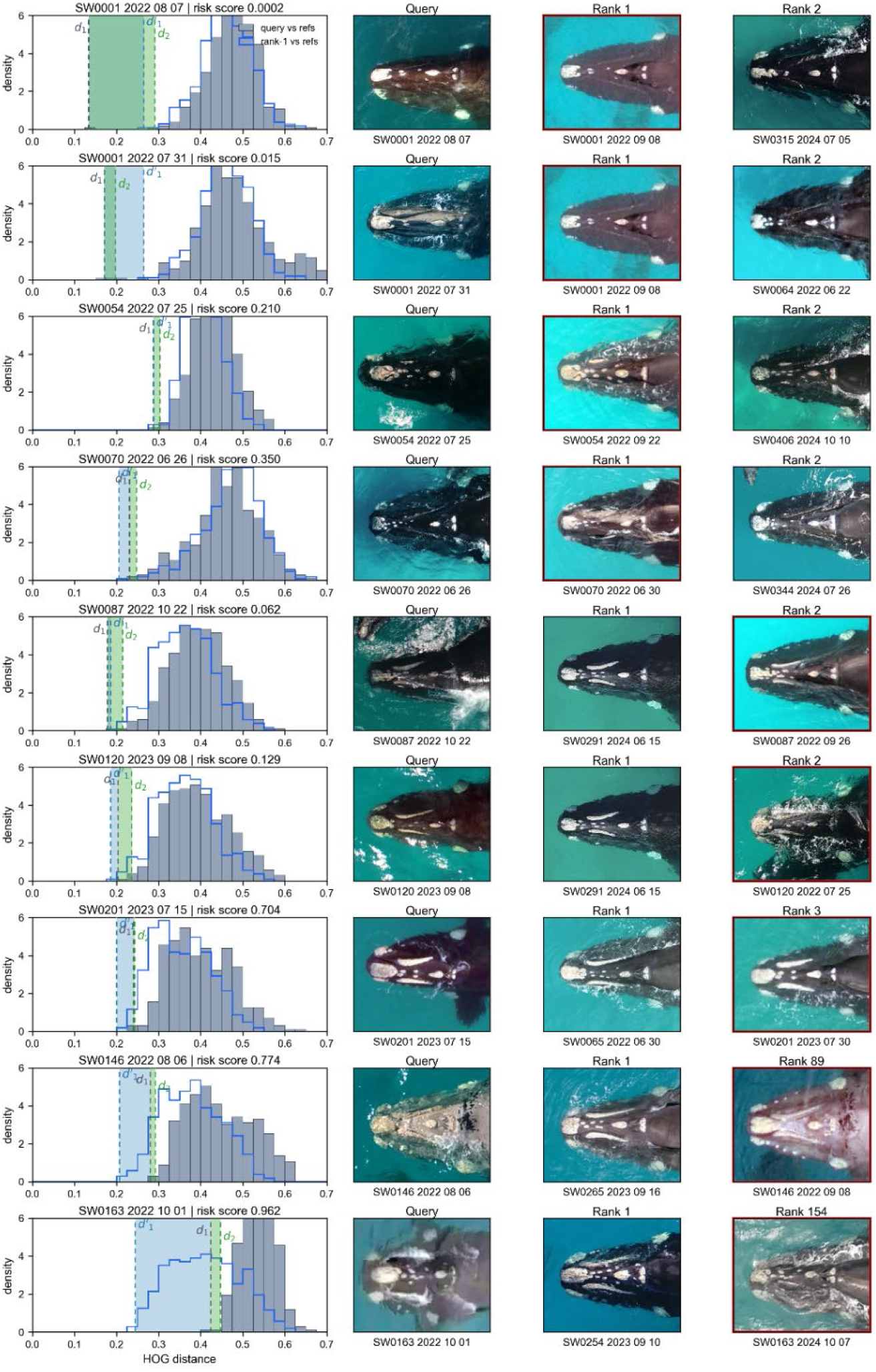
Representative HOG ranking outcomes. Rows show selected validation queries with correct or false rank-1 matches. In the left column, gray histograms show the distribution of HOG distances between the query image and the reference database; the blue outline shows the corresponding distance distribution for the rank-1 reference image. The row title reports the risk score S, i.e. the risk that the first-ranked candidate is incorrect. The gray dashed vertical line marks d_1_. The green dashed vertical line marks d_2_, and the green shaded interval denotes the rank gap g = d_2_ − d_1_. The blue dashed vertical line marks d’_1_, the nearest non-self reference distance of the rank-1 reference image, and the blue shaded interval spans d_1_ and d’_1_, corresponding to the reference-neighbor margin g’ = d’_1_ − d_1_. The image columns show the query, the first-ranked candidate, and either the second-ranked candidate or the known matching whale when the correct match ranked lower than second. Red outlines around image panels mark the known true whale.

**Fig. 3.**
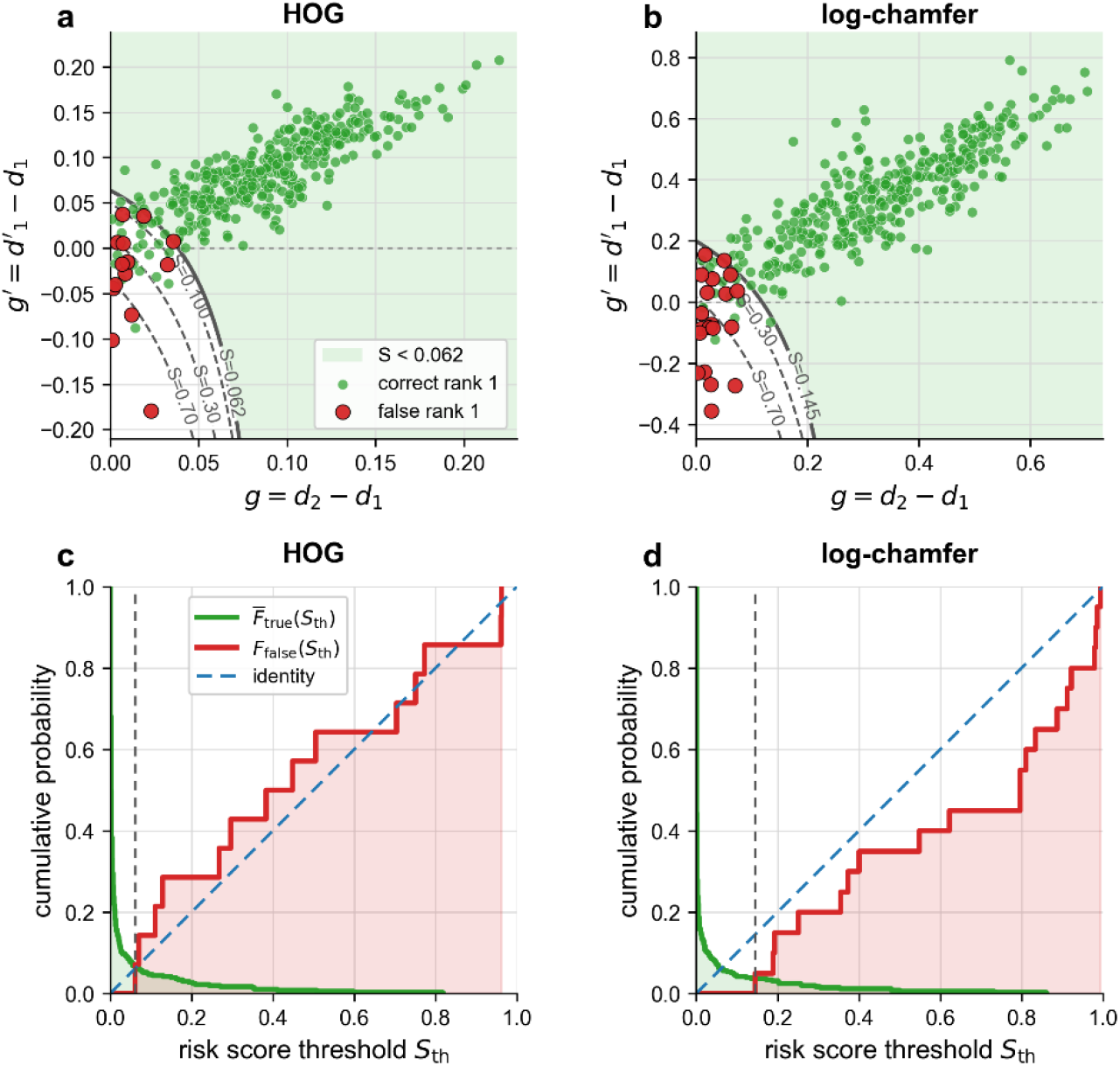
Match-confidence calibration for HOG and log-chamfer ranking. (a,b) Rank-1 score space for HOG (a) and log-chamfer (b). The x-axis shows the rank gap g = d_2_ − d_1_, and the y-axis shows the reference-neighbor margin g’ = d’_1_ − d_1_. Each point corresponds to one validation query; green points are correct rank-1 matches and red points are false rank-1 matches. Gray contour lines show lines of constant risk score S; the pale green region indicates scores below the lowest observed false rank-1 match. (c,d) Empirical cumulative functions for HOG (c) and log-chamfer (d). With increasing risk-score threshold S_th_, the fraction of false matches with S ≤ S_th_ increases, whereas the fraction of correct matches with S ≥ S_th_ decreases. The red curve shows F_false_(S_th_), the green curve shows F̅_true_(S_th_), the blue dashed line is the identity line, and the gray dashed vertical line marks the lowest score observed among false rank-1 matches.

### Initial image registration

Both HOG and chamfer matching used the same image-registration framework. The 750 x 300 pixel ROI crop from the standardized mask was first resized to a 320 x 128 pixel ROI. This 320 x 128 representation was used for both distance calculations and for expressing all translations. To obtain an initial registration seed, the matcher ROI was downsampled further to 80 x 32 pixels. Query templates were generated from this coarse image over rotations from -5 to +5 degrees in 1 degree increments and scale factors from 0.80 to 1.20 in 0.05 increments. Each transformed query template was matched to a padded coarse reference image using OpenCV template matching with the TM_CCOEFF_NORMED similarity metric. The resulting translation was converted back to the 320 x 128 ROI coordinate system and rounded to 4 pixel increments, with translations constrained to +/-60 pixels in that coordinate system. The best OpenCV template match provided one registration seed for each query-reference pair.

### Fine image registration

The initial seed was then refined by greedy local search on the 320 x 128 ROI. For the HOG analysis, each candidate transform was evaluated by recomputing the transformed query HOG descriptor and minimizing the HOG distance to the reference descriptor. For the chamfer analysis, each candidate transform was evaluated by transforming the query and reference edge points and minimizing the symmetric log-chamfer distance. The local search tested neighboring x-y translations, rotations, and scale changes one step away. The fine-search step sizes were 2 pixels for x and y translation in the 320 x 128 ROI coordinate system, 1 degree for rotation, and 0.01 for scale. Transforms were constrained to rotations from -5 to +5 degrees, scales from 0.80 to 1.20, and translations from -60 to +60 pixels. A neighboring transform was accepted only when it lowered the HOG distance or chamfer distance for the respective method, and the search continued until no further improvement was found or 120 iterations were reached. The lower-distance result from the two refined starting transforms was retained for that query-reference pair.

### Registered HOG descriptor distance

For HOG matching, following the general HOG descriptor approach (Dalal & Triggs 2005), callosity edges were first computed on the full standardized binary mask using a 3 x 3 morphological gradient, before cropping to the ROI. The cropped edge image was resized to 320 x 128 pixels using area interpolation, lightly smoothed with a 3 x 3 Gaussian kernel, and protected by a one-pixel zero guard border. This order prevents the ROI boundary from becoming an artificial vertical or horizontal callosity edge. A HOG descriptor was computed using OpenCV’s HOGDescriptor class with a 320 x 128 pixel window, 16 x 16 pixel cells, 32 x 32 pixel blocks, 16 pixel block stride, and nine unsigned orientation bins. This yielded 19 x 7 overlapping blocks, each containing 2 x 2 cells and nine bins, for a 4788-element descriptor. The descriptor was L2-normalized before comparison.

For each registered query-reference comparison, HOG distance was computed as 1 - dot(q, r), where q and r are the normalized HOG descriptors of the transformed query mask and reference mask. This is equivalent to cosine distance for normalized descriptors. For each query image, all reference images were ranked from lowest to highest HOG distance. The best distance and the gap between the best and second-best distances were retained as measures of match quality and ambiguity.

### Registered log-chamfer edge distance

We implemented a modified symmetric chamfer-distance edge matcher using standard OpenCV image-processing operations and the distance-transform approach (Borgefors 1988). Callosity edges were also computed on the full standardized binary mask before ROI cropping, so that ROI borders could not create artificial edge pixels. The ROI edge image was resized to 320 x 128 pixels, and a one-pixel border was suppressed. For each reference mask, a distance image was computed from the inverse edge image using the Euclidean distance transform. Pixel distances were capped at 30 pixels and transformed as log(distance + 1). Edge points were sampled from edge contours, with up to 10 arc-length-spaced points retained per contour, to reduce computational cost while preserving the spatial distribution of the callosity pattern.

For each transform, the query edge points were mapped into the reference coordinate system and sampled from the reference log-distance image. The reference edge points were also inversely mapped into the query coordinate system and sampled from the query log-distance image (Fig. 1). The symmetric log-chamfer distance was the mean of these forward and backward distances, with out-of-bounds points assigned log(31). References were ranked from lowest to highest symmetric log-chamfer distance. As for HOG, the best distance and the best-to-second-best distance gap were retained as quality measures.

### Validation test

To test the image matching performance, we selected 375 query images of 198 individual whales from a catalog with known whale identities. Each query image was compared with all 411 reference images of the catalog, resulting in 154,125 query-reference comparisons per method. A match was counted as successful if the correct whale identity was ranked within a specified candidate list, with top-1, top-5, and top-10 performance reported separately.

### Match confidence

To flag uncertain rank-1 matches after HOG or chamfer ranking for closer visual inspection, we computed the gap *g* between the first- and second-ranked distances, *g = d_2_ – d_1_*. False matches tended to have a small gap, which indicates that the two best-ranked candidates are difficult to distinguish. Although all false matches could be identified by a gap below a threshold value, a considerable fraction of true matches also had gaps below that threshold.

To predict true from false matches with higher discriminatory power, we also computed the smallest distance from the rank-1 reference image to all other reference images, *d’_1_*. If the query image truly matches the rank-1 reference image, then the distribution of query-reference distances should closely resemble the distribution of distances from the rank-1 reference image to all other reference images. In this case, the distances *d_2_* and *d’_1_*, and hence the gap *g’ = d’_1_ – d_1_*, are also expected to be similar to *g*. False matches, by contrast, tended to have a small or negative *g’*, which indicates that the query image and the rank-1 reference image generate distinctly different distance distributions when compared to all other reference images.

From *g* and *g’*, we computed a risk score *S* that best distinguishes correct from false rank-1 matches, based on the results from a validation test. For each query *i*, the risk score *S_i_* was computed with a monotone logistic model:

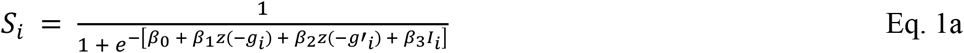

with interaction term *I_i_* = max(0, *z*(−*g_i_*)) max(0, *z*(−*g*′*_i_*)) that is positive only when both *g* and *g’* are small or negative. *z* denotes standardization by the validation-set mean and standard deviation.

The model parameters *β*_1_, *β*_2_, *β*_3_ were fitted from the 375 validation queries by minimizing the L2-regularized norm

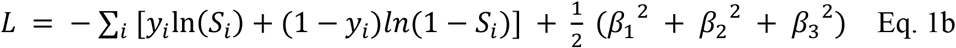

using *y_i_* = 1 for a false rank-1 match and *y_i_* = 0 for a correct rank-1 match. The first term in Eq. 1b penalizes false matches that receive a low risk score, and the second term penalizes correct matches that receive a high risk score. The L2 term discourages unnecessarily large coefficients. The parameters *β*_1_, *β*_2_, *β*_3_ were constrained to positive values to ensure that smaller margins can only increase *S_i_*.

To describe how the risk score separates correct and false rank-1 matches, we computed two empirical cumulative functions from the validation data. For false rank-1 matches, we used the false-match cumulative distribution

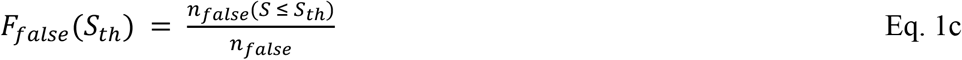

where *n_false_*(*S* ≤ *S_t_*_ℎ_) is the number of false rank-1 validation queries with risk score less than or equal to a threshold score *S_th_*. For correct rank-1 matches, we used the complementary cumulative distribution

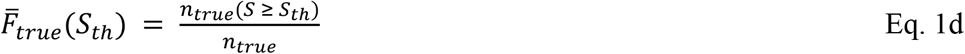

where *n_true_*(*S* ≥ *S_t_*_ℎ_) is the number of correct rank-1 validation queries with risk score greater than or equal to *S_th_*.

### Probability that an unmatched sighting is a new whale

The ranking results can also be used to estimate the probability that a sighting is a new whale when no convincing visual match is found among the top *k* candidates. This requires two quantities. The first is the probability that a recurrent whale is missed within the top *k* ranks, *P*_miss,k_. This value can be estimated from the validation test of *n* validation queries (375 in our case) as

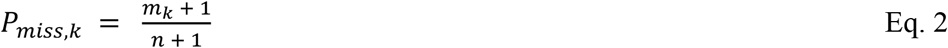

where *m*_k_ is the number of misses among the top *k* ranked candidates. Using the HOG method as an example, the correct whale was not ranked first in 14 cases, thus *m*_1_ = 14; it was ranked below rank 2 in six cases, thus *m*_2_ = 6; and it was ranked below rank 5 in two cases, thus *m*_5_ = 2. The add-one correction in Eq. 2 avoids assigning an over-optimistic empirical miss probability of zero when no top-*k* failures were observed.

The second quantity needed is *P_rec_*, the prior probability that the sighting is recurrent (i.e. the whale is already recorded in the database) before the ranked list is inspected. This prior probability can be obtained from past observations of new or recurrent whales (Fig. 4). If no visually confirmed match is found among the top *k* candidates, Bayes theorem gives:

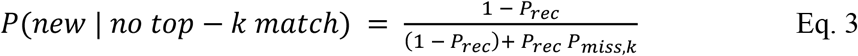

**Fig. 4.**
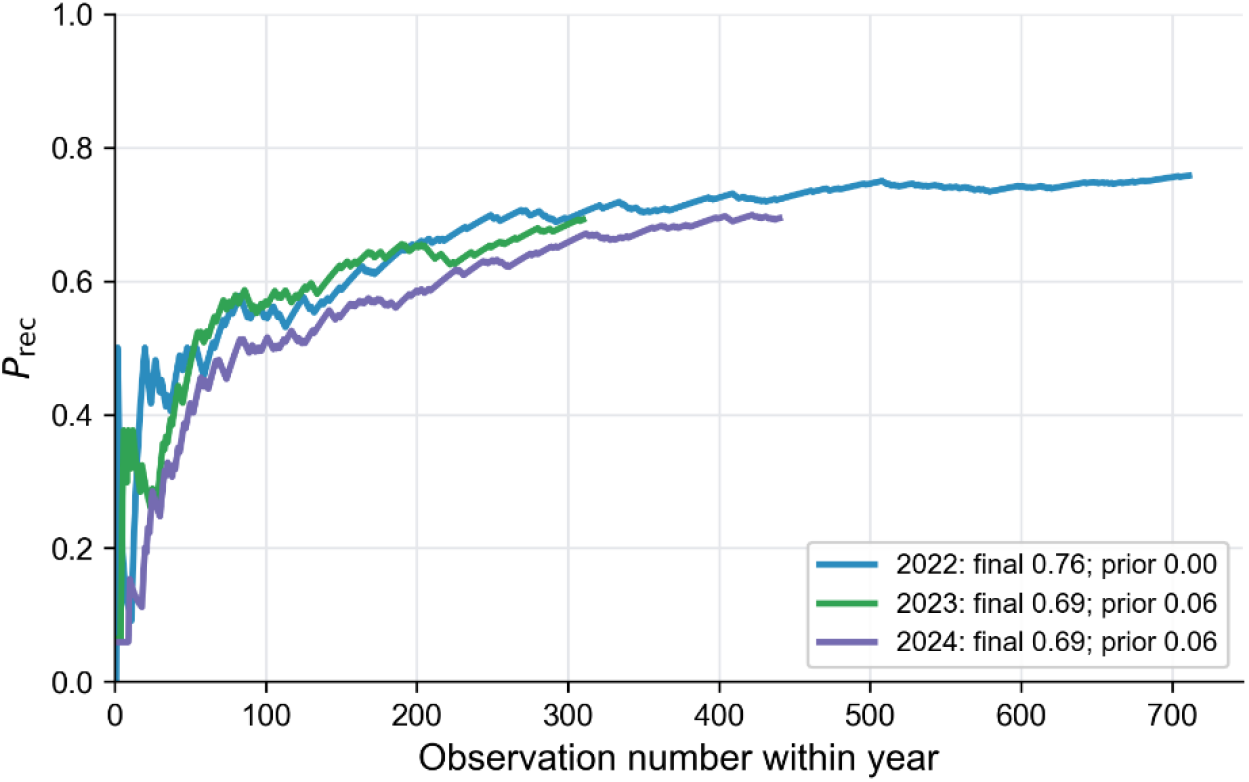
**Running prior probability P_rec_ that a sighting was recurrent**, i.e. that a previously identified whale was observed, separately measured for the 2022, 2023, and 2024 survey seasons. The x-axis is the observation number within each year, starting with 1 for the first sighting of that season. Sightings were ordered by date, time, and database number. Each curve starts at the year-specific prior: 0 in 2022, 0.064 in 2023, and 0.059 in 2024. Thereafter, each already-known whale contributes a value of 1 to the data series, and each first sighting of a new whale contributes the year-specific prior; the plotted value is the cumulative mean of the data series within that year.

**Fig. 5.**
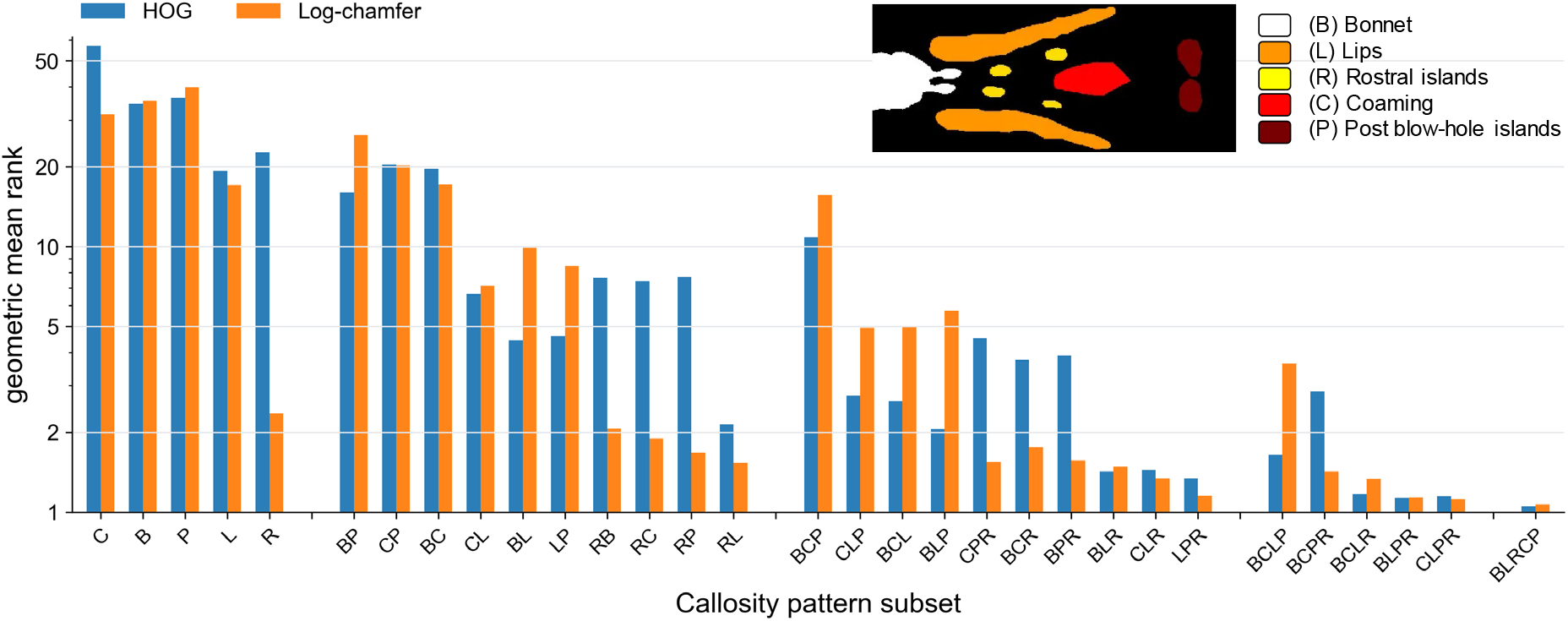
**Method performance for specific callosity pattern subsets**. Selected callosity pattern subsets of candidate whale images from the validation dataset are compared with the 411 reference images using the HOG distance method (blue bars) or the log-chamfer distance method (orange bars), and the geometric mean rank of the correct whales is computed. ‘BLRCP’ refers to the full callosity pattern included, which is the standard image matching procedure; for the meaning of other letter combinations, see inset legend. The inset image shows a representative callosity pattern (SW0400).

Hence, Eq. 3 can be used to stop the visual inspection of ranked match suggestions when the probability of a new whale has increased above a predefined threshold.

### Robustness

To test how the method responded when components of the callosity pattern such as the post blow-hole islands or lips were missing, e.g. due to glare, we systematically removed specific components from the segmented query image masks but not the reference image masks, and computed the HOG and chamfer distance ranking. By systematically adding or removing callosity components, we obtained information on how much each component contributed to the overall matching quality of the method.

### Software implementation and data analysis

Image curation, segmentation review, matching, validation, and data extraction were performed with a custom Whale Server software running in a Docker container on a Linux workstation with Ubuntu 20.04.6 LTS. The workstation was equipped with a 3.6 GHz Intel Core i7-7820X CPU and 125 GiB RAM. The Whale Server used a browser-based single-page frontend implemented in JavaScript, HTML, and CSS, and a Python backend implemented with Flask and served with Waitress. Structured sighting, catalog, and validation data were stored in an SQLite database, whereas image metadata, annotations, masks, and intermediate results were stored as JSON sidecar files and image files. Dedicated Python routines implemented image registration, HOG matching, log-chamfer matching, validation summaries, and figure/data export. All custom software was written with assistance from OpenAI Codex.

## Results

Validation was performed on 375 identified query images. HOG ranked the correct whale first in 361 cases (96.3%), within the top five in 373 cases (99.5%), and within the top ten in 373 cases (99.5%). The 14 HOG misses consisted of eight rank-2 cases, four rank-3 cases, one rank-89 case, and one rank-154 case.

Log-chamfer matching ranked the correct whale first in 355 of 375 cases (94.7%), within the top five in 370 cases (98.7%), and within the top ten in 371 cases (98.9%). The 20 log-chamfer misses consisted of eight rank-2 cases, two rank-3 cases, two rank-4 cases, three rank-5 cases, one rank-9 case, one rank-42 case, one rank-49 case, one rank-121 case, and one rank-284 case.

A simple combined rule selected for each query the first-ranked candidate from the method with the lower risk score *S*. This increased rank-1 performance to 368 of 375 cases (98.1%) and rank-5 performance to 373 cases (99.5%). The seven rank-1 misses were three rank-2 cases, one rank-3 case, one rank-4 case, one rank-89 case, and one rank-154 case.

Fig. 2 illustrates the range of outcomes behind the summary statistics. Correct matches from high-quality query and reference images have a low first distance and a large gap from the next-best candidate. Some correct matches have small gaps because the query image is oblique, the callosity pattern is partially obscured by glare or spray, or differs in contrast from the reference image. False rank-1 matches typically also show a small gap but in addition a shift between the query-reference distance distribution and the rank-1-reference distance distribution (shown by the blue curve in Fig. 2), which is particularly pronounced for the two large-rank mismatches of whales SW0146 and SW0163. The query image for whale SW0146 had poor contrast that makes it difficult to identify the callosity pattern, and the query image for whale SW0163 has been photographed from an oblique angle that causes substantial perspective distortions of the callosity pattern. The chamfer distance distributions of the same selected queries are shown in Fig. S1. Chamfer matching finds the correct identities in all selected cases except for whales SW0146 and SW0163 that were also severely mismatched.

### Match confidence

The risk score converted two warning signs into a single review metric: a small rank gap *g* = *d_2_* − *d_1_* and a small or negative reference-neighbor margin *g’* = *d’_1_* − *d_1_*. For both HOG and log-chamfer, false rank-1 matches clustered at small *g* and small or negative *g’*, whereas most correct rank-1 matches had larger values (Fig. 3A,B). For HOG, the risk score separated false from correct rank-1 matches with an area under the receiver operating characteristic curve (ROC AUC) of 0.979 and an average precision of 0.655. The lowest HOG risk score observed among false rank-1 matches was *S* = 0.062. All 337 validation queries below this score were correct rank-1 matches. For log-chamfer, the risk score separated false from correct rank-1 matches with a ROC AUC of 0.991 and an average precision of 0.861. The lowest log-chamfer risk score observed among false rank-1 matches was *S* = 0.145. All 342 validation queries below this score were correct rank-1 matches.

### Probability that an unmatched sighting corresponds to a new whale

To estimate the probability that an unmatched sighting corresponds to a new whale, we first estimated the prior probability *P*_rec_ that a sighting is recurrent, i.e. the whale has been previously identified and is in the database. At the start of each season, before any new images from that season are reviewed, *P*_rec_ is the fraction of whales in that season that are already known from earlier years. This starting value was 0 in 2022, 7/109 = 0.064 in 2023, and 9/152 = 0.059 in 2024. As each season progresses, the database grows. Each sighting of a whale that has already been recorded contributes a value of 1 to the running estimate. A first sighting of an apparently new whale contributes the starting value for that season. The running mean of this sequence gives the seasonal estimate of *P*_rec_ shown in Fig. 4.

Combining this prior with the validation result allows us to estimate the probability that an unmatched sighting is a new whale. Using HOG as an example, the correct whale was not ranked first in 14 of 375 validation queries and was ranked below the top five in 2 queries. With the add-one correction, *P*_miss,1_ = 15/376 = 0.0399 and *P*_miss,5_ = 3/376 = 0.0080. At the beginning of the 2023 or 2024 season, *P*_rec_ was approximately 0.06. With this prior, failure to accept the first-ranked HOG candidate gives *P*(new | no top-1 match) = 0.997. If visual inspection finds no match among the five top-ranked candidates, this probability increases to *P*(new | no top-5 match) = 0.9995.

By the end of each survey season, *P_rec_* had increased to approximately 0.7. Therefore, *P*(new | no top-1 match) falls to 0.885 for log-chamfer, 0.915 for HOG, and 0.953 for the combination of both methods. If no visual match is found among the top five candidates, *P*(new | no top-5 match) is 0.964 for log-chamfer, and 0.982 for HOG. Accordingly, the search for a match can often be stopped after visually inspecting the first few top-ranked candidates, but the exact stopping threshold should remain conservative and should take image quality, segmentation quality, and distance gaps into account.

### Robustness

To analyze how the method performed when some of the callosity features were missing, e.g. due to glare or head position, we systematically removed specific callosity components from the query image masks of the validation dataset. We then compared the results based on the geometric mean of the rank that was assigned to the correct whale.

With all features included, the geometric mean rank was 1.05 for HOG and 1.10 for log-chamfer. With only one callosity feature included, no single component provided consistently strong performance for both methods, although the rostral islands were the most informative feature for chamfer matching, with a geometric mean rank of 2.56. With two pattern subsets included, rostral islands and lips gave the best results. For three features, rostral islands, lips and post blow-hole islands were best, and for four features, it was the combination of rostral islands, lips, post blow-hole islands, and bonnet. This indicates that the rostral islands are the most informative callosity feature especially for chamfer matching, and the coaming is the least informative callosity feature. Nonetheless, a reliable identification is best achieved by retaining the full callosity pattern whenever possible.

## Discussion

The validation results show that both registered image-matching methods correctly matched most of the query images with reference images from a catalog. HOG was the stronger of the two methods. It ranked the correct whale first in 361 of 375 queries (96.3%) and within the first five candidates in 373 queries (99.5%). The registered symmetric log-chamfer distance placed the correct whale first in 355 queries (94.7%) and within the first five candidates in 370 queries (98.7%). When the first-ranked candidate was selected from the method with the lower risk score, rank-1 performance increased to 368 of 375 queries (98.1%).

Image registration is central to this performance. Even after manual curation, query and reference images differ in head angle, crop placement, scale, and small residual rotations. The OpenCV-seeded registration provides a good initial transform by matching downsampled masks over plausible rotations, scales, and translations. The subsequent fine greedy search then optimizes the method-specific distance at full resolution. This two-stage strategy is faster than a dense exhaustive search while still providing sufficient flexibility to compensate for differences among drone images, such as variation in azimuthal viewing angle.

The two distance measures, HOG and log-chamfer, have complementary strengths. HOG describes the local orientation structure of the callosity mask across overlapping blocks. Log-chamfer matching compares the spatial positions of callosity edges through distance-transform images. It is geometrically intuitive and produces visually interpretable distance maps, but in the current validation it was slower and produced more rank misses than HOG. While the HOG and log-chamfer distance methods performed similarly well when all callosity features were included, the log-chamfer method tended to produce consistently better results when only selected callosity features were used that included the rostral islands. Without the information from the rostral islands, the HOG method tended to perform better. Because the two methods use different representations of the same segmented callosity pattern, using both together improves the matching accuracy.

A useful feature of both approaches is that they do not only return a ranked list of candidate whales, but also indicate when the ranking order is uncertain. The most informative warning sign was the difference in distance, or gap, between the first- and second-ranked candidates. A large gap, together with a similar distance gap in the reference image’s own distance neighborhood, indicates a decisive match. By contrast, a small gap should trigger closer visual review. The method therefore does not only assign an identity but helps prioritize review effort. In the validation test of 375 queries, the correct whale was among the top ten candidates in 99.5% of cases for HOG and 98.9% of cases for log-chamfer. For an experienced matcher, comparing the query image with the first ten reference images requires only 10-20 seconds. If none of the first ten top-ranked reference images is confirmed by visual matching, and if image and segmentation quality are high, the query image likely represents a whale not yet present in the catalog. For HOG, this probability was estimated as 98.2% under a late-season recurrent-sighting prior of *P_rec_* = 0.7.

Compared to state-of-the-art closed-set classifier-based systems, the method presented here provides several advantages. Deep neural-network systems can be highly effective when large, balanced, well-curated catalogs are available, and web platforms can scale them to very large databases. Their main practical costs are the need for substantial labeled training data, population- or viewpoint-specific retraining, and sensitivity to catalog errors and image quality. By contrast, registered HOG and registered log-chamfer matching require only a single curated reference image and segmentation mask per individual, and each new query is compared directly with the current reference database. Adding a new whale requires adding a single curated image and mask, not retraining a classifier. Both methods can be readily implemented on a standard desktop PC. HOG descriptors, template matching, morphological edge extraction, image transformations, and distance transforms are all standard operations available in the Python OpenCV package. Running the matching software does not require specialized GPU hardware or other neural-network infrastructure.

The performance of registered image matching methods depends on accurate segmentation of diagnostic callosities, and poor masks will propagate directly into both descriptor and edge-distance comparisons. Strongly oblique images, incomplete head views, glare, turbidity, sea spray, or unusual posture may also reduce match quality. In this study, the HOG and log-chamfer distance methods were evaluated only on high-quality drone images, most of them taken from a near-nadir perspective, with the whale head viewed approximately from above and with little perspective distortion of the callosity pattern. One of the two large-rank failures in the validation test was caused by an image taken from a more oblique angle. The resulting perspective distortion of the callosity pattern could not be corrected by the current registration procedure, which included only rotation, translation, and scaling. In principle, perspective correction or viewpoint-specific registration could address such cases, but this was outside the scope of the present study.

In conclusion, HOG and log-chamfer matching of drone-derived southern right whale callosity masks provide a practical, training-free approach for individual identification. The empirical risk score derived from rank separation and reference-neighborhood information helps focus manual review on ambiguous cases.

## Acknowledgments

First and foremost, we thank Katy Fannei for her invaluable contributions as a drone pilot, survey scribe and photo-identification matcher, and for her work in maintaining and curating the whale catalogue. We thank all members of the Little White Whale Project for fundraising efforts, helpful discussions, encouragement, and especially acknowledge the Little White Whale Project committee members David Ellett, Lisa-Maree Ellett, Mick Moir, Sue Osborne, and Noel and Robyn Stoney for their generous investment of time and tireless effort in managing all logistical matters. A special and heartfelt thank you goes to Robyn and Noel Stoney for their extraordinary generosity in providing housing to our research team and for their advice in all matters. We are grateful to the following individuals and organizations for allowing us access to their private properties to reach critical aggregation sites: Wayne Batson of Australia Civil Haulage, Lisa and Travis Bone, Nick and Anna Gorman, Bill and Anna Cagnana, Quaalup Homestead, the Hassell family, Trevor of the Pallinup and the certified legendary Randall. We thank Graeme Drew for providing free accommodation during extended survey expeditions in Bremer Bay. We acknowledge the efforts of the following volunteers who contributed to data entry and ID curation: Nicola Adcock, Olivia Anderson, Liz Bosh, Holly Butterworth, Andrew and Deanna Davenport, Kalagan Driver, Sam and Lilly Edwards, Celeste Grim, Alex Grim, Michele Hannibaldsen, Shendell Hay, Robin Heudebert, Chloe Hodkinson, Kirry Joyce, Heqi Li, Sinead McMahon, Marta Mrasic, Skyler Schmitter, Anne Schmock, Ellie, Zoe and Alby Slatter, Renee Sumich, Paul Trela, Sari Tuffi, Renae Van Noort, Lily Warren, and Holly Williams. We thank Kate Sprogis and Chandra Salgado-Kent for their valuable discussions and insights. We are especially grateful to David Campbell Jr. and Sr. of Campbell Transport, whose donation of the 4-wheel drive vehicle for surveys and all associated fuel costs made this entire research endeavor possible. Without their generosity, none of the survey work could have happened. Technical support was provided by Nik Callow and David Tunbridge from the University of Western Australia. We acknowledge the Traditional Custodians of the land and sea country on which this research was conducted, and pay our respects to Elders past, present and emerging.

## Author Contributions

Ben Fabry: Conceptualization, Methodology, Software, Formal analysis, Data curation, Visualization, Writing - original draft, Writing - review and editing. Greta Jacobs: Data curation, Validation, Formal analysis, Writing - review and editing. Johannes Bartl: Methodology, Software, Writing - review and editing. Max Fabry: Conceptualization, Investigation, Formal analysis, Resources, Data curation, Project administration, Funding acquisition, Writing - review and editing. All authors reviewed and approved the submitted manuscript.

## Data Accessibility

The complete whale image catalog, field metadata, and curated identification records used in this study are not publicly available as a complete package. The authors are willing to share selected images for specific scientific purposes, including curated reference images and segmented callosity masks. The authors can also compare external southern right whale images against the study catalog to support collaborative identification, cross-catalog matching, or independent validation.

## Ethics Statement

Animal ethics approval was granted through the Animal Ethics Committee of the University of Western Australia (project 2021/ET000236).

## Supplementary Information

**Supplementary Fig. S1.**
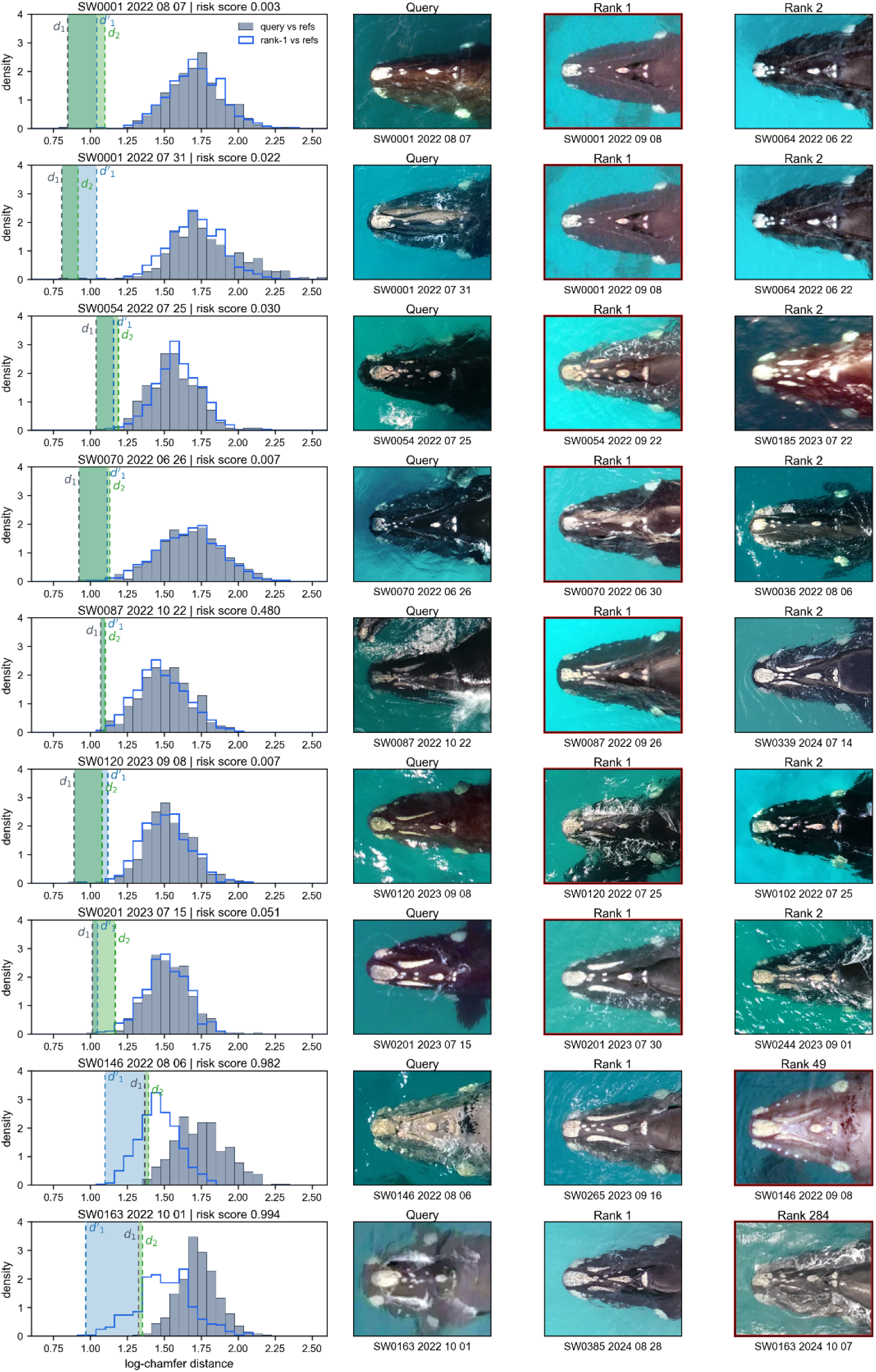
Representative chamfer ranking outcomes. Gray histograms show the distribution of chamfer distances between the query image and the reference database; the blue outline shows the corresponding distance distribution for the rank-1 reference image. The row title reports the risk score S. The gray dashed vertical line marks d_1_. The green dashed vertical line marks d_2_, and the green shaded interval denotes the rank gap g = d_2_ − d_1_. The blue dashed vertical line marks d’_1_, the nearest non-self reference distance of the rank-1 reference image, and the blue shaded interval spans d_1_ and d’_1_, corresponding to the reference-neighbor margin g’ = d’_1_ − d_1_. The image columns show the query, the first-ranked candidate, and either the second-ranked candidate or the known matching whale when the correct match ranked lower than second. Red outlines around image panels mark the known true whale.

